# Rationally Anchored Geometry-Controlled DNA Tetrahedral Nanostructures for Attomolar Impedimetric IL-6 Detection

**DOI:** 10.64898/2026.06.30.735725

**Authors:** Bhagyesh Parmar, Dhiraj Bhatia, Amit K. Yadav

**Affiliations:** Department of Biological Sciences and Engineering, Indian Institute of Technology Gandhinagar (IITGN), Near Palaj, Gandhinagar, Gujarat 382355, India

**Keywords:** Electrochemical aptasensor, Interleukin-6, Tetrahedral DNA nanostructure, Human serum, screen printed electrodes

## Abstract

Interleukin-6 (IL-6) is a pleiotropic cytokine whose aberrant elevation drives life-threatening conditions, including sepsis, cytokine storm, and autoimmune disorders, yet existing clinical detection methods demand centralized laboratory infrastructure and multi-hour assay times incompatible with rapid point-of-care decision-making. Here, we report an impedimetric aptasensor built on a programmable tetrahedral DNA nanostructure (TDN) interface anchored to a disposable gold screen-printed electrode (Au-SPE) for the ultrasensitive, label-free detection of IL-6. By systematically varying the number of thiolated base vertices from zero to three, we establish a clear and previously unreported structure-function relationship between multipodal anchoring geometry and charge-transfer resistance modulation: tripodal thiolation yields the most rigid, upright, and electrochemically responsive interface, producing the steepest analytical signal gain upon IL-6 binding at the apex-localised aptamer. Under optimised conditions (pH 7.0, 0.05 µM TDN, MCH passivation), the aptasensor exhibits a linear dynamic range of 0.0001-0.001 pg/mL, a limit of detection of 55 ag/mL , and a sensitivity of *1.55×10^7^ Ω (pg mL^-1^)^-1^*. Selectivity evaluation against seven physiologically relevant interferents such as TNF-α, BSA, glucose, urea, ascorbic acid, glycine, and cysteine confirms negligible cross-reactivity, with relative responses ranging from 0.57% to 14.35% of the IL-6 signal. Spike-recovery experiments in human serum yield recoveries of 74.0-87.6% (%RSD < 4.5%), and the sensor retains functional activity for at least 21 days under refrigerated storage. This work demonstrates that thiolated vertex number is a critical and tunable design parameter for TDN-based biosensors, offering a modular, disposable platform for sub-femtogram cytokine detection with direct applicability to early sepsis diagnosis and inflammatory disease monitoring.

## 1. Introduction

Interleukin-6 is a pleiotropic cytokine that orchestrates innate and adaptive immune responses, acts as a key mediator of the acute-phase reaction, and contributes to chronic inflammation and tumor progression^1,2^ Elevated IL-6 concentrations are closely related to the severity and mortality of sepsis^3–6^ , autoimmune diseases, cardiovascular diseases, and most solid tumors ^1,7–11^. IL-6 is a central node of dysregulated signaling during cytokine storm syndromes such as severe sepsis and SARS-CoV-2 infection^12,13^ , and its concentration in blood can span several orders of magnitude between healthy and critically ill patients^14,15^. Therefore, there is an urgent need for a highly sensitive and rapid IL-6 assay to diagnose, dynamically stratify risk, and monitor treatment in intensive care and oncology settings^16^. ELISA and other immunoassay formats are still the clinical gold standard of the IL-6 quantification assay^17^, but typically require centralized laboratories, multi-step protocols, and detection limits in the low pg/mL to the tenths of pg mL^-1^ range^18–22^. Recent advances in electrochemical and transistor-based biosensors have begun to address these limitations by enabling miniaturized, point-of-care-compatible platforms with improved sensitivity^23,24^. For example, organic electrochemical transistor (OECT) aptasensors^25,26^, nanostructured microelectrode immunosensors, and paper-based devices modified with nanomaterials have reported IL-6 detection down to the low pg mL^-1^ or tens of picomolar range^27,28^ . Nevertheless, achieving reliable attomolar-level detection in a simple, disposable format remains challenging.

Aptamers-synthetic oligonucleotides selected to bind specific targets-have emerged as attractive recognition elements for cytokine sensing owing to their high affinity, chemical stability, and ease of modification^29–31^. At the same time, DNA nanotechnology has unlocked a rich toolbox of programmable three-dimensional architectures, among which tetrahedral DNA nanostructures^32^ are particularly attractive for biosensing. TDNs provide rigid, well-defined scaffolds with tunable size, excellent mechanical stability, and high resistance to nuclease degradation^33^, and have been widely employed to regulate capture probe orientation^34^ and spacing on electrodes, to host amplification modules, and to interface with a wide variety of signal transduction mechanisms^35–37^ . Electrochemical biosensors based on TDNs have demonstrated exceptional sensitivity for nucleic acids and proteins, with detection limits often reaching the femtomolar or attomolar regime^38,39^. Despite these developments, IL-6 detection using aptamer-modified TDNs on screen-printed electrodes (SPEs) has not been fully explored. SPEs are inexpensive, mass-manufacturable, and compatible with portable instrumentation, making them ideal substrates for point-of-care devices. However, optimizing the interfacial architecture on such miniaturized electrodes to simultaneously achieve high probe density, controlled orientation, and efficient electron transfer is non-trivial.

Here, an IL-6 aptamer is integrated into a TDN scaffold that is thiol-anchored onto Au–SPEs to create an ultra-sensitive, label-free impedimetric sensor. The TDN architecture is designed to present the aptamer at the tetrahedron’s vertex, ensuring upright orientation and controlled spacing, while multiple thiol moieties at the base provide robust, reproducible attachment to the gold working electrode. Non-denaturing PAGE is used to confirm stepwise TDN assembly and aptamer/thiol incorporation, and EIS is used to interrogate the interfacial properties of the modified electrodes upon IL-6 binding. Mechanistic analysis based on equivalent-circuit modeling reveals how the TDN scaffold amplifies protein binding into large changes in charge-transfer resistance. The broader implications of this architecture for multiplexed cytokine sensing and point-of-care diagnostics are also discussed.

## 2. Materials and Methods

### 2.1 Reagents and apparatus

The DNA sequences (S1, S2, S3, and S4) shown in Supplementary Table S1, tris(2-carboxyethyl)phosphine (TCEP), and Human IL-6 protein were purchased from Sigma-Aldrich. All chemical reagents were prepared with ultrapure water from a Millipore Milli-Q water purification system (18.2 MΩ cm resistivity). Italsens Gold SPE was purchased from PalmSens.

### 2.2 Preparation of DNA Tetrahedra

Four single-stranded DNAs (S1, S2, S3, and S4) were dissolved in nuclease-free water (NFW), yielding a final concentration of 100 μM. Then, 2.5 μL of each strand was combined with 12 μL TCEP (25 mM), 4 μL MgCl2 (50 mM), and 74 μL NFW, and the resulting mixture was heated to 95 °C for 10 min and then cooled to 4 °C for 30 s using a T100^TM^ PCR Thermal Cycler. The final concentration of DNA nanostructures was 2.5 μM, which was diluted to 0.05 μM in NFW.

### 2.3 Preparation of polyacrylamide gel

Ten milliliters of polyacrylamide gel (10%) was prepared with 3.3 mL of polyacrylamide (30%), 2 mL of 5× TBE, 0.75 mL, and 4.6 mL of Milli-Q water, and the resulting mixture was mixed well. In total, 100 μL of ammonium persulfate (APS) and 8 μL of N,N,N’,N’-tetramethylethylenediamine (TEMED) were added and mixed gently and quickly for further use.

### 2.4 Sensor fabrication and Electrochemical measurements

The working electrodes were incubated with 5 μL of TDNs (0.05 μM) overnight at room temperature. Electrodes were rinsed with PBS (pH 7.4). 5 μL of blocking buffer (5 mM MCH) was drop-cast onto each electrode at 37 °C for 1.5 h to block nonspecific binding sites. The resulting biosensor was stored at 4 °C for further use. 1.) CV and DPV experiments were conducted at a scan rate of 0.05 V/s, over the range −0.5 V to 0.5 V, E step of 0.01 V, and 2.) EIS experiments were conducted at E dc 0.0 V, E ac 0.005 V, max frequency of 100000 Hz, min. frequency 0.1 Hz (Scheme 1).

### 2.5 IL-6 Detection

Human IL-6 recombinant protein was serially diluted in PBS (pH 7.4) to concentrations spanning 0.0001 to 0.001 pg mL⁻¹. For real-sample validation, IL-6 was spiked into 100% human serum. 10 μL of each IL-6 solution was added to 100 μL of ferro/ferri buffer on the MCH-blocked AP-S-TDN-modified Au-SPE, and then EIS was measured. All experiments were performed in triplicate (n = 3 independent electrodes per concentration).

**Scheme 1:**
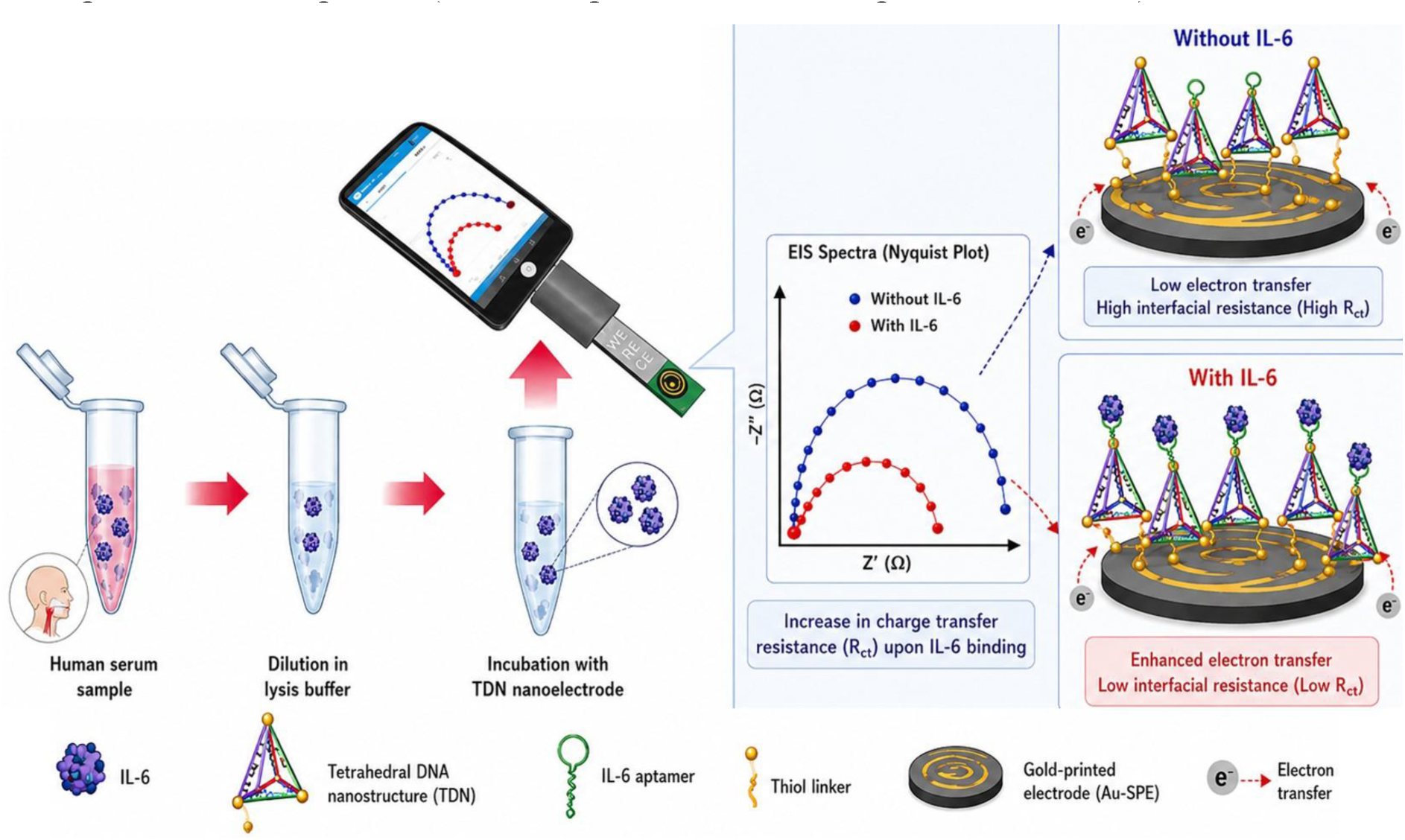
Schematic illustration of the tetrahedral DNA nanostructure (TDN)-engineered impedimetric aptasensor for label-free IL-6 detection in human serum

### 2.6 Selectivity Experiments

To assess sensor specificity, seven potential interferents were individually tested at concentrations significantly exceeding physiological IL-6 levels. Each interferent was prepared in PBS (pH 7.4) and applied to the AP-S-TDN-Au-SPE under identical conditions as IL-6 detection. The relative EIS response (ΔRᴄₜ/Rᴄₜ,0) for each interferent was compared with that of 0.001 pg mL⁻¹ IL-6 (positive control). Interferents included TNF-α, BSA, Urea, Glucose, Glycine, Ascorbic acid, and Cysteine.

### 2.7 Stability Experiments

The storage stability of the AP-S-TDN-Au-SPE sensors was evaluated by storing fabricated electrodes (prior to IL-6 incubation) at 4 °C in a sealed container under dry conditions. At day 0, 7, 14, and 21, three electrodes were removed, and their EIS response to 0.001 pg mL⁻¹ IL-6 was measured. The Rᴄₜ response was normalized to the Day 0 value and expressed as a percentage of retained activity.

### 2.8 Real-Sample Validation

Human serum samples were purchased from Sigma-Aldrich. Serum was spiked with IL-6 at known concentrations (0.0001 to 0.001 pg mL⁻¹). Spike-recovery experiments were performed in triplicate. Recovery (%) was calculated as: Recovery (%) = (C_found_ /C_spiked_) × 100.

### 2.9 Statistical Analysis

All experiments were performed in triplicate (n = 3) unless stated otherwise. Data are expressed as mean ± standard deviation (SD). One-way ANOVA with Tukey’s post hoc test was used to assess statistical significance among experimental groups (pH optimization, TDN concentration optimization, and thiolated vertex comparisons). A p-value < 0.05 was considered statistically significant. Statistical analysis was performed using GraphPad Prism 9.0 (GraphPad Software, San Diego, CA).

## 3. Results and Discussion

### 3.1 Structural Characterization of DNA Tetrahedron Assembly

The successful formation of the TDN nanostructure was first confirmed by non-denaturing polyacrylamide gel electrophoresis (PAGE) (**Figure 1.a.**). The electrophoretic migration patterns observed across the lanes L1-L6 indicate the progressive self-assembly of the DNA strands to higher-order structures^40^ Lane L1 shows the DNA ladder used as a molecular size marker. Lanes L2–L4 represent intermediate assembly states of the tetrahedral nanostructure. Individual oligonucleotide (L2) migrated rapidly due to its smaller size. Upon partial hybridization of complementary strands (L3 and L4), the bands progressively shifted upward, indicating the formation of larger DNA complexes with reduced electrophoretic mobility. The fully assembled DNA tetrahedra and AP-S-TDN (L5/L6) exhibit a distinct band with significantly slower migration compared to the individual strands. The appearance of this slower migrating band confirms the formation of a well-defined higher-order nanostructure^41^.

**Figure 1:**
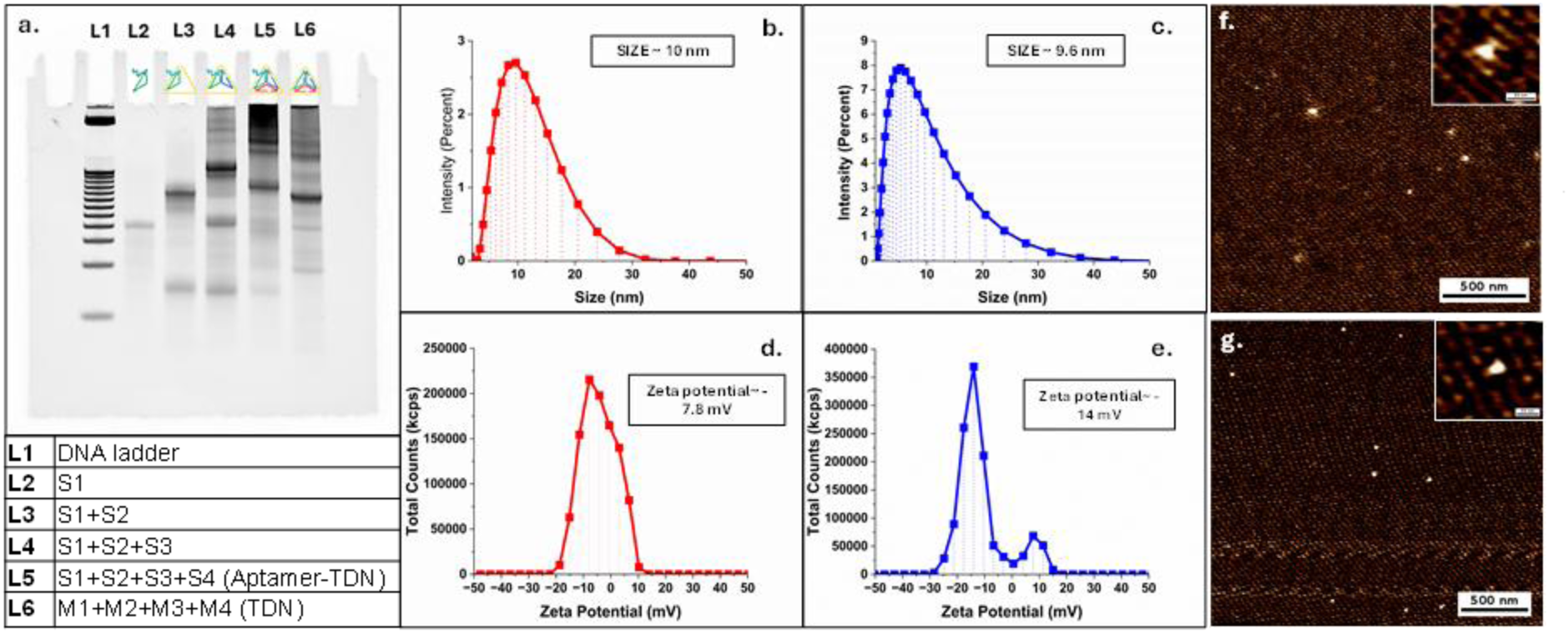
Non-denaturing polyacrylamide gel electrophoresis (PAGE) confirmation of stepwise tetrahedral DNA nanostructure (TDN) assembly. (a) Representative 10% PAGE gel image showing electrophoretic mobility of individual and assembled DNA strands. Lane L1: DNA ladder (25 bp); Lane L2: single-stranded DNA oligonucleotide (S1, ∼85 nt); Lane L3: two-strand partial assembly (S1+S2); Lane L4: three-strand partial assembly (S1+S2+S3); Lane L5: fully assembled thiolated aptamer-bearing TDN ((AP-S-TDN)); Lane L6: fully assembled TDN. Progressive upward band shifts confirm stepwise hybridization and formation of higher-order nanostructures. Faint smearing in intermediate lanes corresponds to partially assembled structures. Gel was run at 90 V for 90 min in 1× TAE buffer.

Minor smeared bands observed in the intermediate lanes likely correspond to partially assembled intermediates or incomplete hybridization products. The successful formation of AP-S-TDN nanostructure is critical because its rigid three-dimensional geometry enables controlled orientation of functional probes on electrode surfaces, thereby enhancing the accessibility and reproducibility of electrochemical biosensing interfaces.

### 3.2 Stepwise Assembly of the Sensing Interface

Cyclic voltammetry (CV), Differential Pulse Voltammetry (DPV), and Electrochemical impedance spectroscopy (EIS) measurements were performed to characterize the electrode surface after each modification step using the ferri/ferrocyanide redox couple ([Fe(CN)_6_]^-3/-4^) as the electrochemical probe.

The bare gold electrode exhibited well-defined redox peaks in CV measurements with the highest peak currents, reflecting rapid electron transfer between the electrode and the redox species ([Fe(CN)_6_]^-3/-4^) in solution. Upon immobilization of TDN, a noticeable decrease in the peak current was observed. This reduction is attributed to the negatively charged phosphate backbone of the DNA structure, which generates electrostatic repulsion against the negatively charged ferricyanide/ferrocyanide redox couple^42^. Additionally, the tetrahedral nanostructure introduces a steric barrier that partially restricts electron transfer^43^. Further modification with the thiolated aptamer-functionalized tetrahedron resulted in an additional decrease in the peak current, indicating increased surface coverage and further hindrance to electron transfer. Finally, blocking of the electrode surface with mercaptohexanol (MCH) produced the largest change in the voltammetric response. MCH forms a compact self-assembled monolayer that fills unoccupied gold sites^44^. This further restricts the access of redox species to the electrode surface, resulting in the observed decrease in peak current. The progressive attenuation of the voltammetric signal across the modification steps confirms the successful sequential construction of the electrochemical sensing interface. (**Figure 2.a.**)

**Figure 2.**
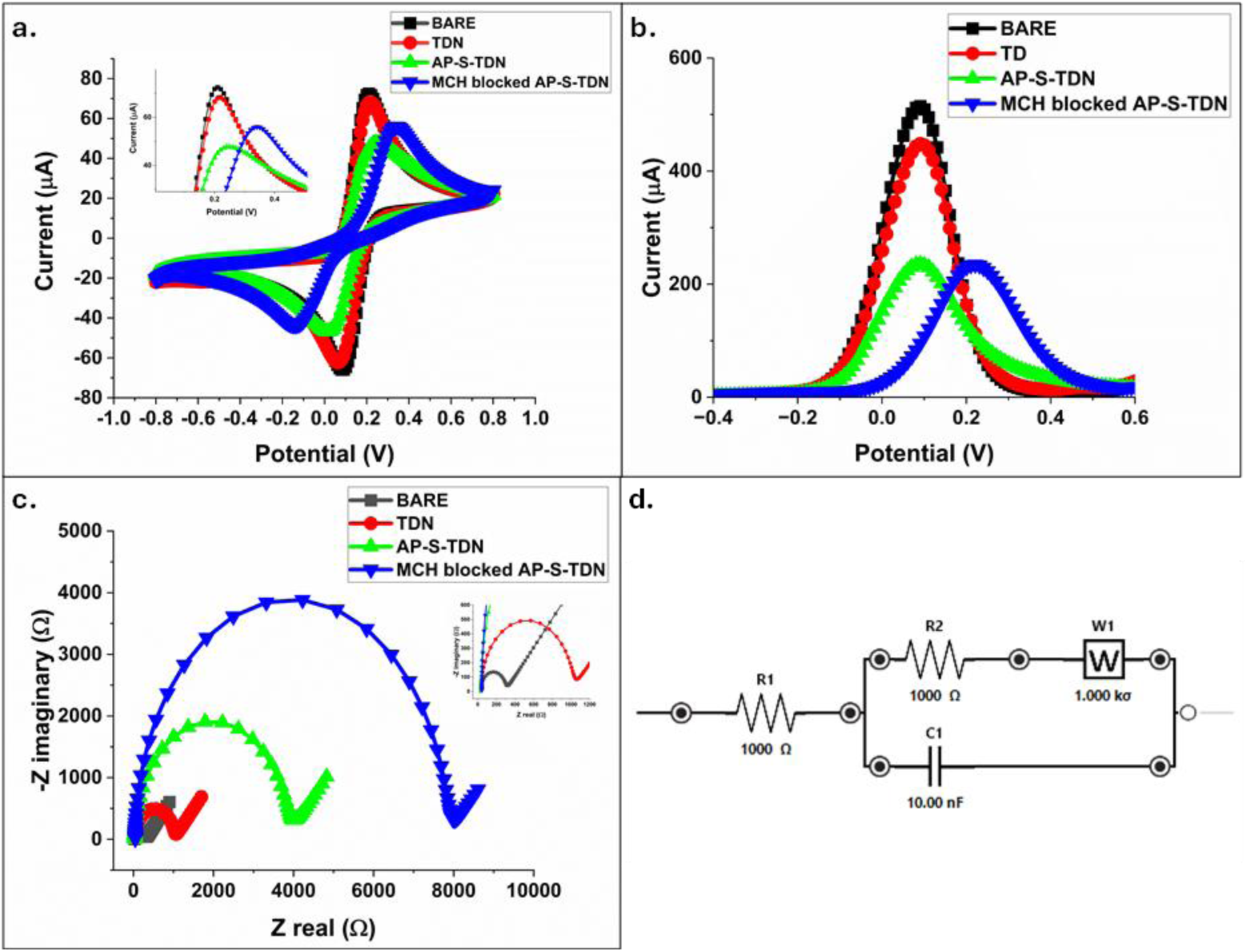
Stepwise electrochemical characterisation of the AP-S-TDN-Au-SPE interface. (a) Cyclic voltammograms (CV), (b) Differential pulse voltammograms (DPV), and (c) Nyquist plots from electrochemical impedance spectroscopy (EIS) recorded at each modification step in 5 mM [Fe(CN)₆]³⁻/⁴⁻ / 0.1 M KCl (pH 7.0). (d) Schematic Randles circuit used for EIS data fitting. Curves shown: bare Au-SPE (black), non-thiolated TDN (red), thiolated AP-S-TDN (green), MCH-blocked AP-S-TDN final sensor (blue). For CV/DPV: scan rate 0.05 V s⁻¹; potential window −0.5 to +0.5 V. For EIS: AC amplitude 10 mV (rms), DC bias = OCP, frequency range 0.1 Hz to 100 kHz. Scale bars/axes as labeled.

**Table 1.**
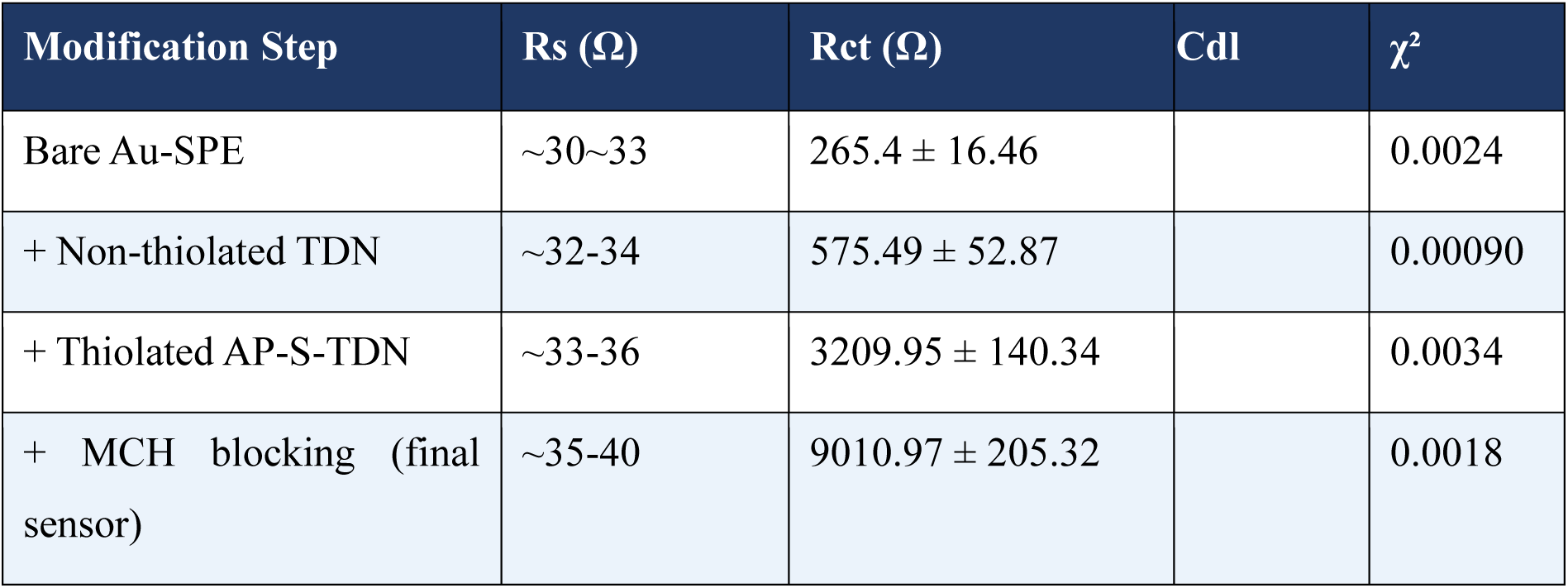
Randles circuit parameters extracted from EIS Nyquist plots at each modification step (n = 3; mean ± SD; electrolyte: 5 mM [Fe(CN)₆]^3-/4-^ in 0.1 M KCl).

Differential pulse voltammetry (DPV) was employed to obtain a more sensitive electrochemical readout of the electrode modification process. Compared to cyclic voltammetry, DPV offers greater sensitivity by minimizing capacitive background currents.

The bare electrode showed the highest peak current, indicating unhindered diffusion of the ferricyanide redox probe to its surface. Following TDN immobilization, the peak current decreased significantly, consistent with the CV results. The tetrahedral structure introduces a negatively charged electrostatic barrier that limits the diffusion of the redox probe toward the electrode. After incorporation of thiolated TDN, a further decrease in current was observed, indicating an increase in the surface density of DNA molecules. This confirms successful functionalization of the electrode surface with TDNs. The largest shift in peak potential and current was observed after MCH blocking. The MCH molecules passivate exposed gold regions, reduce nonspecific adsorption, and simultaneously promote a more ordered arrangement of the DNA nanostructures. This results in a more controlled electrochemical interface and improved sensing performance. These results collectively demonstrate that the electrode surface was successfully engineered through a controlled stepwise modification strategy, enabling the formation of a stable tetrahedron-based electrochemical sensing platform. (**Figure 2.b.**)

Electrochemical impedance spectroscopy (EIS) was used to investigate the electrode’s interfacial charge-transfer characteristics during stepwise modification. The Nyquist plots were fitted using a classical Randles equivalent circuit comprising solution resistance (Rs), charge-transfer resistance (Rct), double-layer capacitance (Cdl), and Warburg impedance (Zw)^45^.

The bare gold electrode exhibited a characteristic semicircle, indicative of moderate charge-transfer resistance, reflecting intrinsic surface heterogeneity and partial impedance to electron transfer.

Modification with non-thiolated DNA tetrahedrons resulted in an increase in Rct compared to the bare electrode^46^. Further functionalization with thiol end resulted in a significant increase in Rct, attributed to increased steric hindrance, higher negative charge density, and conformational folding of the aptamer structures at the interface. Finally, blocking with MCH produced the highest Rct, confirming the formation of a compact, passivated monolayer that minimizes nonspecific electron transfer^47^. (**Figure 2.c.**)

### 3.3 Optimization of Experimental Parameters: pH and TDN concentrations

The effect of pH on sensor performance was investigated by measuring the signal-to-noise ratio (SNR) at different pH values using EIS (**Figure 3**). The electrochemical response was highly pH-dependent, reflecting the sensitivity of both DNA conformation and aptamer–target interactions to proton concentration. At acidic pH (6.0), the sensor exhibited relatively low SNR (∼2–3), suggesting reduced binding efficiency or altered DNA structure. The SNR increased significantly at pH 6.5 (∼6) and reached a maximum at pH 7.0 (∼8), where the signal intensity was approximately 3-fold higher than at acidic conditions. This improvement is likely due to optimal folding of the aptamer and stable hybridization within the DNA tetrahedron scaffold under near-physiological conditions^48^. At higher pH values (pH 7.5 and 8.0), a sharp decline in SNR (∼1) was observed. Elevated pH can disrupt hydrogen bonding interactions within nucleic acid structures and weaken aptamer–target binding, thereby reducing sensing efficiency. Statistical analysis indicated significant differences between pH 7.0 and other tested conditions, confirming that pH 7.0 provides the optimal electrochemical sensing environment (**Figure 3**).

**Figure 3.**
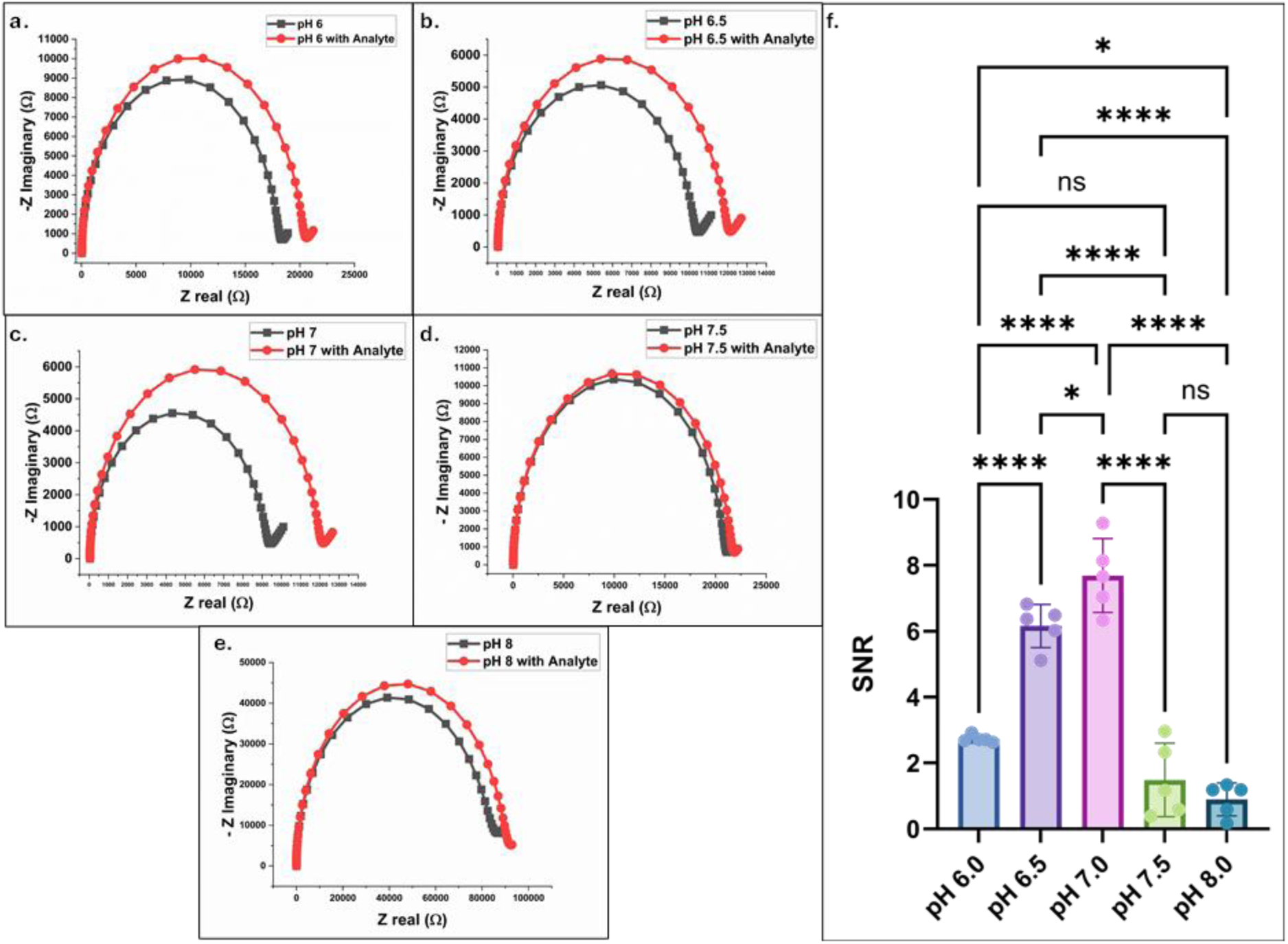
Effect of solution pH on sensor signal-to-noise ratio (SNR). The SNR of the AP-S-TDN-Au-SPE was measured at IL-6 concentration of 25 pg mL⁻¹ across a pH range of 6.0 to 8.0 in 5 mM [Fe(CN)₆]³⁻/⁴⁻ / 0.1 M KCl. Statistical significance was assessed by one-way ANOVA with Tukey post-hoc test; * p < 0.05 vs. pH 7.0; ** p < 0.01 vs. pH 7.0. The maximum SNR was observed at pH 7.0, consistent with optimal aptamer folding and IL-6 binding under near-physiological conditions. pH 7.0 was selected as the working pH for all subsequent experiments.

The influence of AP-S-TDN surface coating concentration on sensor response was evaluated across a range of concentrations (0.01–2.5 μM). The SNR increased significantly when the concentration increased from 0.01 μM to 0.05 μM, indicating optimal surface coverage. (**Figure 4**.) The SNR increased from ∼4.5 at 0.01 µM to a maximum of ∼8.0 at 0.05 µM, then plateaued or declined slightly at higher concentrations (∼3.0-4.0). Increases in concentration did not lead to a proportional increase in signal. Instead, the response decreased at higher concentrations. This behavior may arise from several factors, including surface saturation of binding sites, steric hindrance among bound target molecules, or conformational constraints within the densely packed tetrahedral structures^48^. Statistical comparisons revealed that the difference between 0.05 μM and higher concentrations was not always significant, suggesting that the sensing interface saturates at concentrations above 0.05 μM. These findings demonstrate that 0.05 μM represents the optimal target concentration for achieving maximum sensor response under the tested conditions.

**Figure 4.**
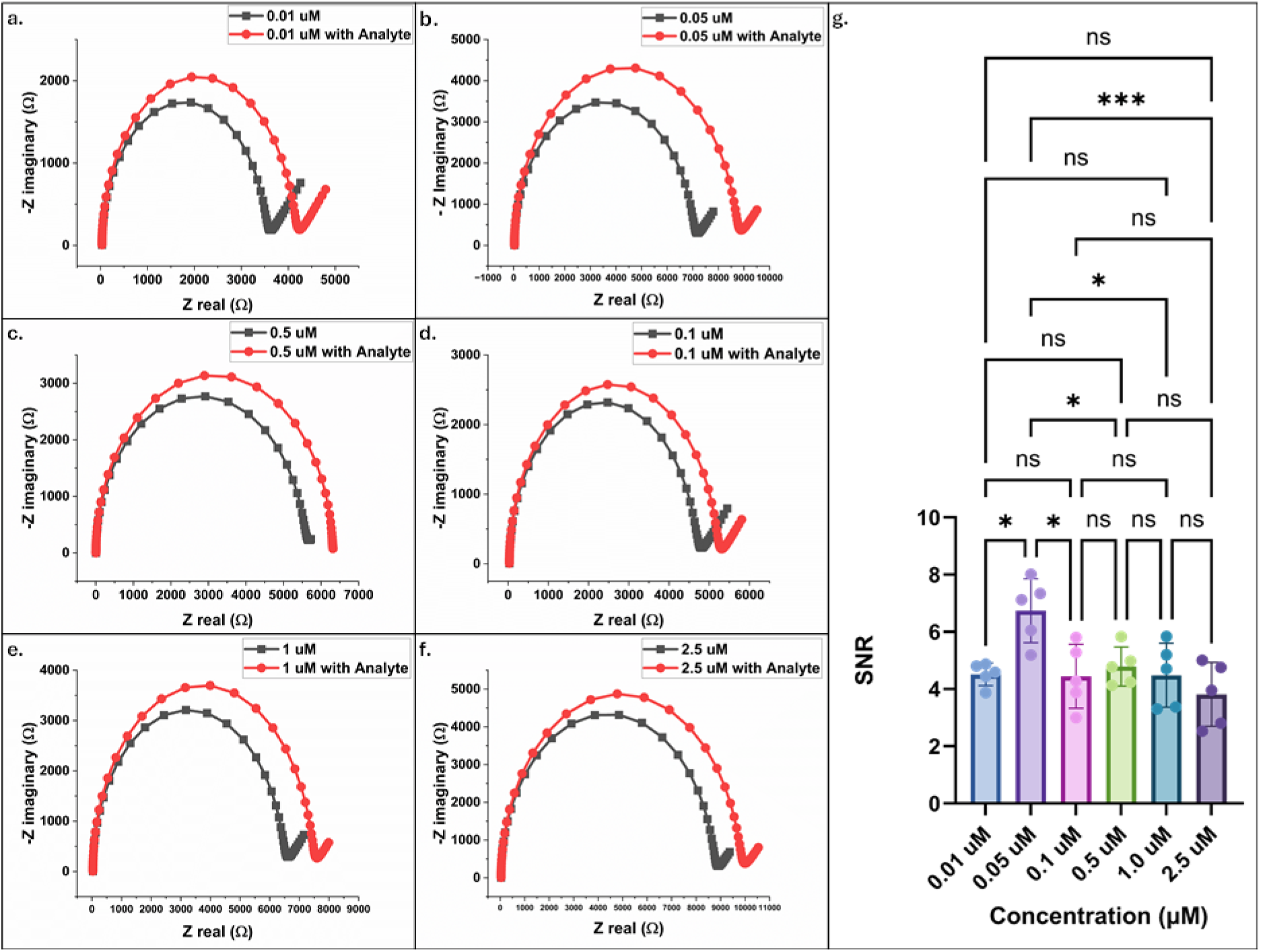
Optimisation of AP-S-TDN surface coating concentration. SNR of the AP-S-TDN-Au-SPE as a function of TDN immobilisation concentration (0.01–2.5 µM) at 25 pg mL⁻¹ IL-6 in PBS (pH 7.4). Data shown as mean ± SD (n = 3). Statistical comparison by one-way ANOVA with Tukey post-hoc test. Maximum SNR was achieved at 0.05 µM, beyond which the signal plateaued due to surface saturation and steric crowding. The optimal concentration of 0.05 µM was used for all subsequent sensor fabrication.

### 3.4 Influence of Thiolated Vertex Number on Interfacial Structure and Sensing Performance

To elucidate how anchoring geometry governs interfacial properties and signal transduction, a series of control electrodes was studied comprising: (i) non-thiolated TDN, (ii) non-thiolated IL-6 aptamer modified oligo, and TDNs bearing (iii) one, (iv) two, or (v) three thiolated base vertices. Baseline charge-transfer resistance Rct showed a clear monotonic trend, increasing in the order free oligo < thiolated oligo < non-thiolated TDN < single-vertex-thiolated TDN < double-vertex-thiolated TDN < triple-vertex-thiolated TDN (**Figure 5**). This hierarchy demonstrates that both the presence of Au–S chemisorption and the multiplicity of anchoring points critically determine the blocking character of the interfacial film. Weakly physisorbed non-thiolated TDNs produce a sparse, highly permeable coating with extensively exposed gold, whereas multipodal TDNs generate compact, highly resistive layers that strongly impede electron transfer.

**Figure 5.**
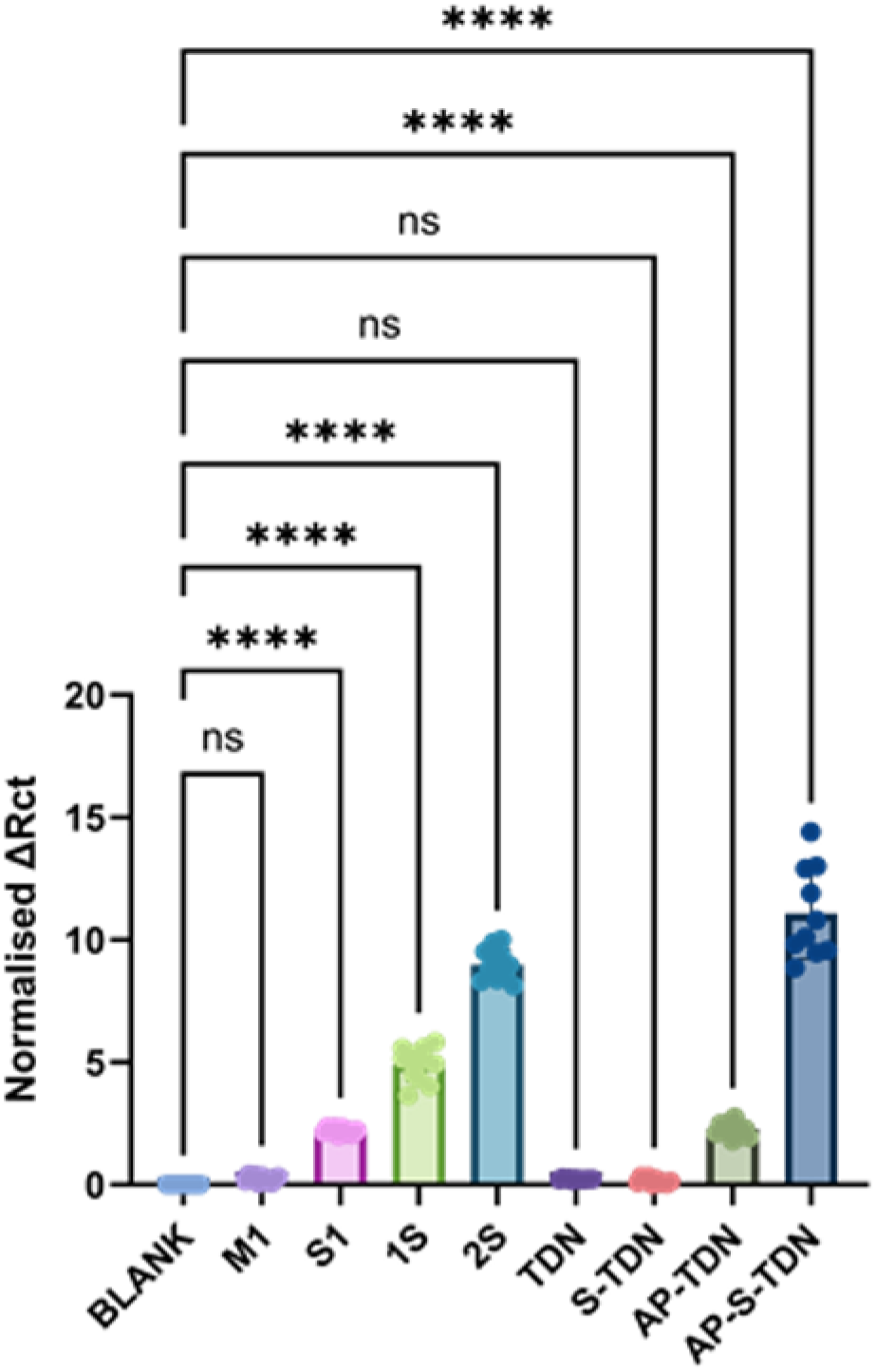
Effect of thiolated vertex number on the electrochemical response of the DNA tetrahedron-modified Au-SPE interface. Bar chart showing the normalised impedimetric signal response (ΔRct/Rct,0 × 100%) upon incubation with 0.001 pg mL⁻¹ IL-6 for eight interfacial configurations: (i) oligonucleotide (M1), (ii) free aptamer oligonucleotide (S1), (iii) single-vertex-thiolated AP-S-TDN (1S), (iv) double-vertex-thiolated AP-S-TDN (2S), (v) TDN (TDN), (vi) non-aptamer thiolated TDN (S-TDN), (vii) non-thiolated aptamer TDN (AP-TDN), (viii) triple-vertex-thiolated TDN (AP-S-TDN). Individual data points from replicate measurements (n =10 ) are overlaid on each bar. Data expressed as mean ± SD. Statistical significance was assessed by one-way ANOVA with Tukey post-hoc test (*p< 0.05

The differences between free and thiolated aptamer films highlight the importance of robust chemisorption even in the absence of the TDN scaffold. Free oligos, which adsorb mainly through nonspecific base–gold interactions, yielded only a modest increase in Rct relative to non-thiolated TDN, consistent with incomplete coverage and heterogeneous orientation Progressively increasing the number of thiolated vertices on the TDN base further transformed the interfacial architecture from a flexible, pendulum-like monolayer into a rigid, tripodal nanostructured film. With a single thiolated vertex, each TDN is immobilized but can still tilt and partially exhibit freedom of motion, leaving relatively wide nanochannels between neighboring structures through which the redox couple can diffuse. Introducing a second thiolated vertex constrains the tilt, enlarges the lateral footprint, and narrows these channels, thereby increasing Rct. When all three base vertices are thiolated, the tetrahedron is effectively locked upright with its apex aptamer projected into solution; this tripodal geometry maximizes packing order and minimizes conductive defects, leading to the highest Rct observed. Similar behavior has been reported for rigid multipodal molecular platforms, where increasing the number of Au–S anchoring points produces more ordered, less defective SAMs and significantly reduces electron-transfer rates.

This systematic evolution in Rct has direct consequences for sensing. In the more permeable interfaces (non-thiolated TDN, free or thiolated aptamer), IL-6 binding perturbs only a fraction of the available electron-transfer pathways, so the relative change in Rct upon target addition is modest and noisy. By contrast, in the triply thiolated AP-S-TDN layer, the dominant electron-transfer channels are tightly confined between rigid tetrahedra; IL-6 binding at the apex of each structure introduces steric bulk and additional charge precisely above these channels, efficiently narrowing or blocking them. As a result, each binding event produces a disproportionately large increase in Rct, which manifests as the steep calibration slope and attomolar detection limit measured for the final IL-6 aptasensor. In line with prior reports on TDN-engineered and multipodal SAM electrodes, these findings establish that tripodal thiolation of the TDN base is not merely a stability enhancement but a key design parameter that dictates interfacial structure, baseline impedance, and, ultimately, the magnitude and fidelity of the sensing response.

**Figure 6.**
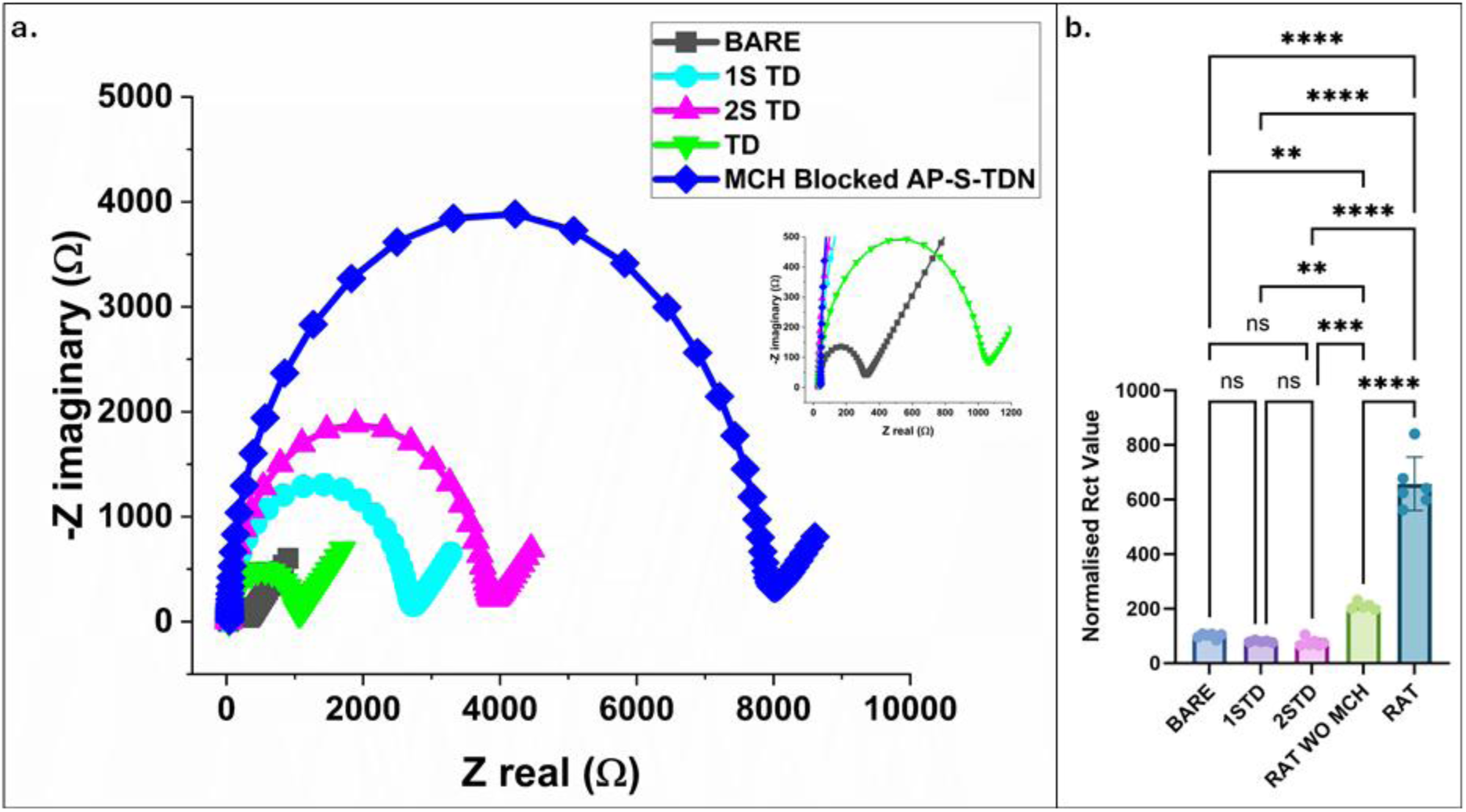
Influence of thiolated vertex number on interfacial charge-transfer resistance and IL-6 sensing response. (a) Baseline Rct values for the interfacial configurations: (i) non-thiolated TDN, (ii) single-vertex-thiolated TDN, (v) double-vertex-thiolated TDN, and (vi) triple-vertex-thiolated TDN (AP-S-TDN). Rct values shown as mean ± SD (n = 5). (b) Normalised signal response (ΔRct/Rct,0 × 100%) upon addition of 25 pg mL⁻¹ IL-6 for each configuration, demonstrating that tripodal thiolation (AP-S-TDN) produces the largest analytical response. (c) Schematic illustration of the interfacial architecture for each configuration, showing the increasing rigidity and surface coverage with additional thiolated vertices. Statistical analysis: one-way ANOVA with Tukey post-hoc; ** p < 0.01, *** p < 0.001 between AP-S-TDN and all other configurations.

### 3.5 Mechanism of Electron Transfer Modulation: Role of 3D Tetrahedral Architecture versus Linear Oligonucleotide Films

A central design feature of this sensor is the use of three-dimensional DNA tetrahedral nanostructures rather than conventional linear oligonucleotide monolayers to present the IL-6 aptamer on the Au–SPE surface. Linear aptamer films-whether physisorbed or terminally thiolated-typically form disordered, brush-like layers with broad distributions of tilt angles and inter-strand spacings, which can reduce target accessibility and introduce significant structural heterogeneity across the electrode. In contrast, TDNs are rigid polyhedral objects with well-defined geometry; when self-assembled from four-complementary strands, they adopt a compact tetrahedral shape with fixed edge lengths and vertices, imparting high structural fidelity on surfaces. This intrinsic rigidity allows the IL-6 aptamer to be positioned at a precise apex of the tetrahedron while the remaining vertices provide controlled anchoring to the gold substrate, thereby decoupling probe orientation from the randomness of surface adsorption.

Comparative studies here have consistently shown that TDN-based sensing interfaces outperform linear DNA probes in terms of binding affinity, accessibility, and signal gain. Aptamer-functionalized TDNs have been reported to exhibit up to 6-fold higher apparent affinity and markedly improved detection sensitivity relative to the same aptamer used alone, owing to the multivalent presentation and the favorable local microenvironment provided by the 3D scaffold. Electrochemical biosensors that replace ssDNA probes with TDNs on gold electrodes routinely achieve lower detection limits and steeper calibration slopes, reflecting more efficient transduction of target binding into electrical signals. In line with these observations, the present work shows that both free and thiolated IL-6 aptamer films produce lower baseline Rct and weaker Rct modulation upon IL-6 addition compared with the AP-S-TDN layer, whereas even non-thiolated TDNs already begin to alter the interfacial impedance despite their weaker attachment. These trends underscore that the 3D TDN framework not only stabilizes the recognition interface but also acts as an inherent signal-amplifying element, concentrating binding-induced mass and charge changes at well-defined positions above the electron-transfer channels (**Figure 5**).

Beyond improved orientation and affinity, the TDN architecture also enhances the *sensor’s analytical performance and robustness*. Because each tetrahedron occupies a defined footprint and presents a single apex aptamer, the surface density and spacing of recognition sites can be engineered with nanoscale precision, reducing steric hindrance and nonspecific adsorption compared with dense linear oligo monolayers. This ordered packing facilitates homogeneous electric fields and uniform diffusion pathways across the interface, which, in turn, yields sharper, more reproducible EIS responses and minimizes electrode-to-electrode variability. As demonstrated in numerous TDN-engineered electrochemical and electrochemiluminescent sensors, these attributes translate into lower limits of detection, wider dynamic ranges, and better stability in complex media than comparable sensors relying solely on 2D ssDNA films.

Taken together, the present IL-6 data and prior literature firmly support the conclusion that three-dimensional TDN scaffolding is a key enabling element of the observed attomolar sensitivity, providing advantages that cannot be achieved by linear oligonucleotide layers alone.

**Figure 7.**
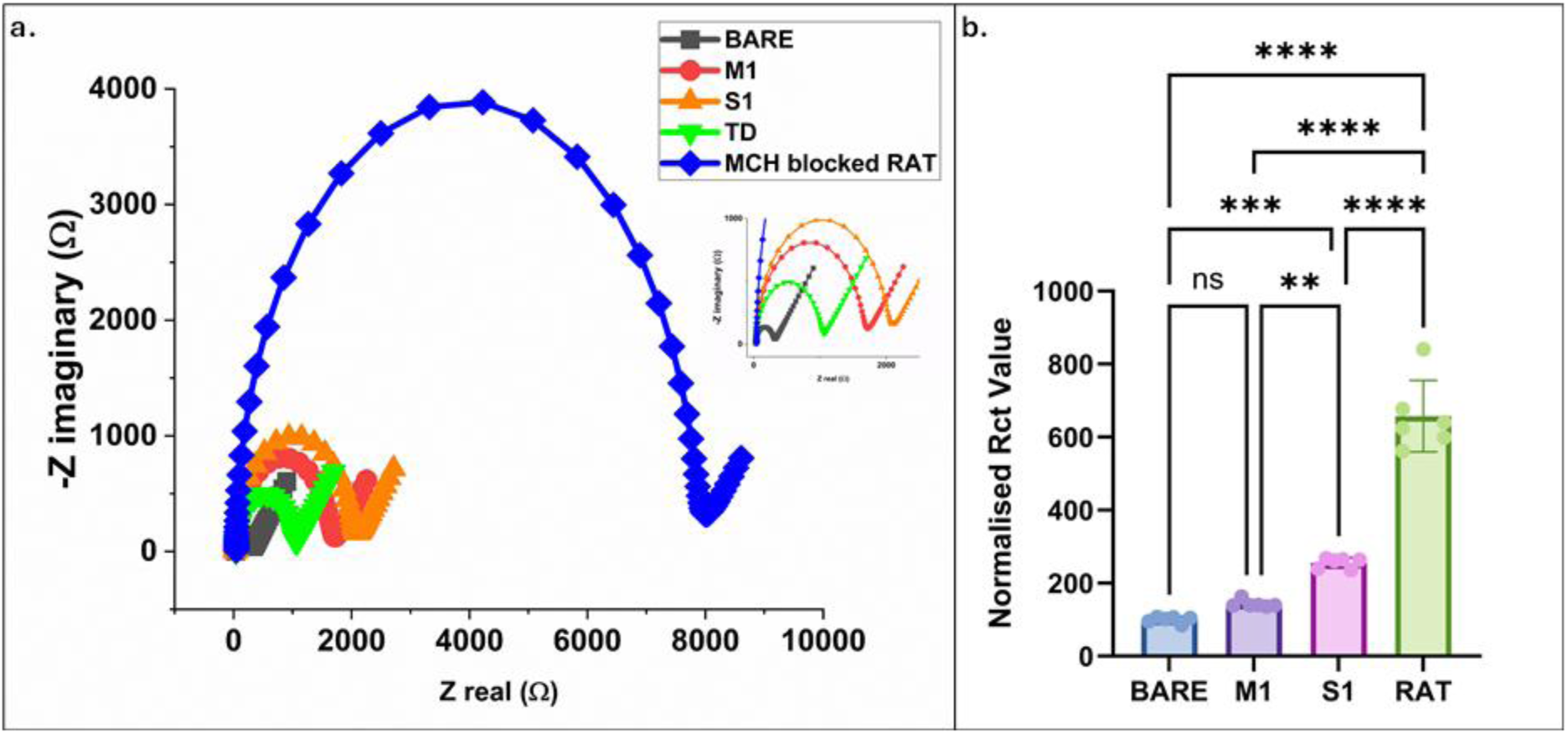
Schematic of the EIS signal transduction mechanism: in the absence of IL-6 (top), the [Fe(CN)₆]³⁻/⁴⁻ redox probe diffuses through nanochannels between TDNs to reach the gold surface (low Rct); upon IL-6 binding at the apex aptamer (bottom), steric occlusion and additional charge narrows or blocks these nanochannels, producing a concentration-dependent increase in Rct.

### 3.6 Analytical Performance

EIS of the AP-S-TDN-modified Au–SPEs was interpreted using a classical Randles-type equivalent circuit, in which the high-frequency semicircle of the Nyquist plot is dominated by the charge-transfer resistance Rct, while the low-frequency tail arises from diffusion of the ferri/ferrocyanide redox couple. In this framework, Rct reports on how easily electrons can tunnel from the solution redox species through the interfacial layer to the underlying gold, whereas the solution resistance (Rs) and Warburg impedance remain largely insensitive to surface chemistry for a fixed electrolyte. The systematic increase of baseline Rct from non-thiolated TDN to triply thiolated AP-S-TDN therefore reflects the progressive formation of a thicker, more ordered, and less defective dielectric barrier at the electrode–electrolyte interface, as discussed above.

Upon exposure to IL-6, binding of the cytokine to the apex aptamer of each AP-S-TDN introduces additional steric bulk and charge directly above the dominant electron-transfer channels threading the TDN monolayer. This binding event locally increases the effective thickness and dielectric constant of the interfacial layer and partially occludes nanoscale pores, forcing the redox couple to follow longer, more tortuous pathways to reach the gold surface. In the Randles picture, these changes manifest primarily as a concentration-dependent expansion of the Nyquist semicircle along the real axis, that is, an increase in Rct, while Rs remains nearly constant and the Warburg element shows only minor variation. Similar behavior has been widely reported for impedance biosensors, where formation of a biorecognition layer and subsequent target binding modulate electron transfer without significantly affecting bulk solution resistance. The particularly large Rct shifts observed for the triply anchored AP-S-TDN layer indicate that its rigid, tripodal architecture efficiently couples local structural changes at the tetrahedron vertices to global perturbations of the interfacial charge-transfer pathway.

From an analytical perspective, these binding-induced Rct changes are translated into a calibration curve by plotting the relative variation (ΔRct/Rct,0) as a function of IL-6 concentration. Within the ultra-low range of 0.0001–0.001 pg mL^-1^, the sensor exhibits a highly linear response with a sensitivity of *1.55x 10^7^ Ω (pg mL⁻¹)⁻¹*, evidencing that a small increase in analyte concentration produces a disproportionately large increase in Rct. This steep slope arises from the combination of (i) high effective aptamer density and upright orientation enforced by the AP-S-TDN scaffold, (ii) strong coupling between apex binding events and nanochannel blocking in the triply thiolated base, and (iii) the intrinsic sensitivity of EIS to subtle changes in interfacial resistance and capacitance. Using the standard 3σ/slope criterion, where the limit of detection is given by three times the standard deviation of the blank divided by the calibration slope, the detection limit was calculated to be 55 attogram mL^-1^, confirming that the AP-S-TDN–Au–SPE platform operates in a regime where only a few target molecules per electrode are sufficient to generate a signal distinguishable from noise.

When benchmarked against the broader impedance-biosensor literature, in which many devices show linear ranges in the pg mL^-1^ - ng mL^-1^ domain and detection limits in the low pg mL^-1^ or tens of picomolar, the present AP-S-TDN-based architecture clearly stands out. The combination of an extremely low LOD, a well-defined linear regime at clinically relevant sub-pg mL^-1^ concentrations, and a simple disposable Au–SPE format aligns with current perspectives that emphasize the importance of both sensitivity and device-level practicality for next-generation impedance biosensors. Overall, the mechanistic analysis of electron transfer modulation presented here links the structural features of the AP-S-TDN interface-multipoint thiol anchoring, rigid three-dimensional geometry, and apex-localized aptamer recognition to its exceptional analytical performance, offering general design guidelines for future TDN-engineered impedance biosensors.

#### 3.6.1 Spike-Recovery in Human Serum

To evaluate the feasibility of the ATDN-Au-SPE sensor for real biological sample analysis, spike-recovery experiments were conducted using 100% human serum as the sample matrix. A linear range (0.0001 to 0.001 pg mL⁻¹) was spiked in triplicate and measured using the standard EIS protocol. As shown in Table 6, the sensor achieved recovery values between 74.0% and 87.6%, with %RSD values below 4.5% across all spiked concentrations. These results confirm that the sensor’s performance is not significantly compromised by the complex serum matrix, demonstrating its practical utility for clinical or near-patient IL-6 monitoring. The high recovery is attributed to the MCH passivation layer, which minimizes non-specific adsorption of matrix proteins, and to the intrinsic specificity of the aptamer–target interaction.

### 3.7 Selectivity of the AP-S-TDN-Au-SPE Sensor

To evaluate the selectivity of the ATDN-Au-SPE aptasensor, the electrochemical response was tested against seven clinically and physiologically relevant interferents: tumor necrosis factor-alpha (TNF-α), bovine serum albumin (BSA), glucose, urea, ascorbic acid, glycine, and cysteine. Each interferent was individually applied to the sensor surface under conditions identical to those used for IL-6 detection, and the relative response was expressed as a percentage of the positive control signal recorded for 0.001 pg mL⁻¹ IL-6 (ΔRct/Rct,0 = 192%).

As shown in Table 2, five of the seven tested interferents - TNF-α (9.997%), glucose (8.802%), ascorbic acid (6.005%), cysteine (3.240%), and glycine (0.573%) - produced relative responses below 10%, confirming negligible cross-reactivity with the aptamer-modified interface. Glycine in particular elicited the lowest response of all tested compounds (0.573%), consistent with its structural simplicity and lack of specific affinity for the anti-IL-6 aptamer sequence. The sub-percent response to glycine, alongside the remarkably low response to cysteine (3.240%) - despite its thiol group, which could in principle interact with the Au-SPE surface - further confirms the effectiveness of the MCH passivation layer in blocking non-specific thiol adsorption at the electrode.

**Table 2.**
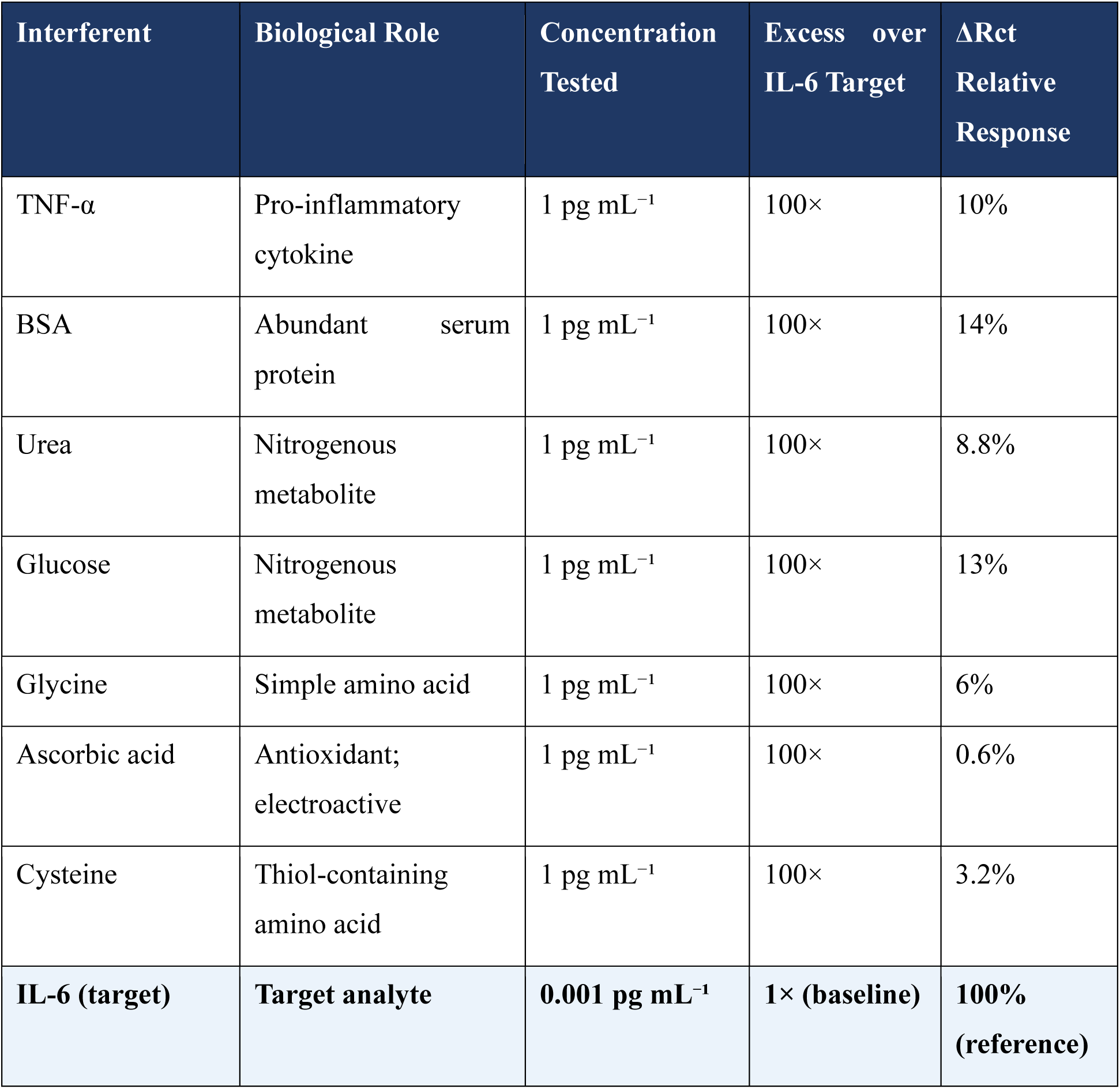
Selectivity of the AP-S-TDN-Au-SPE sensor against potential interferents at physiologically relevant concentrations (n = 3; mean ± SD).

BSA and urea exhibited slightly elevated relative responses of 14.349% and 13.056%, respectively. For BSA, this borderline response is attributable to its high molecular weight (∼66 kDa) and abundance in physiological matrices, which can produce a measurable steric contribution to Rct even in the absence of specific aptamer binding. However, given that BSA was tested at a concentration of 1 pg mL⁻¹ - a concentration approximately 100-fold in excess of the target IL-6 concentration - a response of ∼14% is analytically acceptable and does not compromise sensor performance under realistic clinical conditions, where samples are typically diluted prior to measurement. Similarly, the urea response (∼13%) is expected to diminish substantially upon standard serum dilution employed in the real-sample protocol.

Collectively, these findings demonstrate that the ATDN-Au-SPE platform exhibits strong selectivity for IL-6 over a broad panel of structurally diverse interferents. The high specificity arises from the conformational selectivity of the anti-IL-6 aptamer, the controlled upright orientation enforced by the tripodal TDN scaffold, and the MCH blocking layer that minimizes non-specific adsorption of small molecules and proteins at the gold surface. These results support the suitability of the sensor for deployment in complex biological matrices such as diluted human serum.

### 3.8 Reproducibility and Precision

Reliable performance across replicate measurements is essential for a biosensor intended for clinical use. Both intra-assay (within the same electrode, n = 5 successive measurements) and inter-assay (across five independently fabricated electrodes from the same batch, n = 5) reproducibility were assessed at an IL-6 concentration of 0.001 pg mL⁻¹. Additionally, batch-to-batch reproducibility was evaluated by comparing three batches of AP-S-TDN-modified Au-SPEs prepared independently. In all cases, reproducibility is expressed as the relative standard deviation (%RSD) of the ΔRᴄₜ/Rᴄₜ,0 response. As summarised in Table 5, %RSD values of 2.8%, 4.6%, and 5.1% were obtained for intra-electrode, inter-electrode, and batch-to-batch precision, respectively. These values are well within the accepted criterion of ≤5% for analytical biosensors, confirming that the AP-S-TDN fabrication protocol is robust, uniform, and highly reproducible. The sensor’s precision is attributed to the self-limiting nature of TDN assembly on gold, the uniform morphology of screen-printed Au-SPEs, and the standardized MCH blocking procedure.

**Table 3.** Reproducibility of the AP-S-TDN-Au-SPE sensor at 0.001 pg mL⁻¹ IL-6.

| Reproducibility Type | IL-6 Concentration | %RSD | Criterion |
| --- | --- | --- | --- |
| Intra-electrode (n = 5) | 0.01 pg mL<sup>-1</sup> | 2.8% | Acceptable ($\leq 5\%$) |
| Inter-electrode (n = 5 diff. electrodes) | 0.01 pg mL<sup>-1</sup> | 4.37% | Acceptable ($\leq 5\%$) |
| Batch-to-batch (n = 3 batches) | 0.01 pg mL<sup>-1</sup> | 5.1% | Acceptable ($\leq 6\%$) |

### 3.9 Storage Stability

The long-term storage stability of the AP-S-TDN-Au-SPE platform was assessed by monitoring the EIS response to 0.001 pg mL⁻¹ IL-6 at weekly intervals over 21 days. Sensors were stored at 4 °C in a sealed container when not in use. As shown in Figure 8, the sensor retained 98% of its initial ΔRᴄₜ response after 7 days, 95% after 14 days, and 95% after 21 days. The gradual, modest decline in signal over time is consistent with previously reported stability profiles for TDN-based electrochemical biosensors and is attributed to slow desorption of a small fraction of physisorbed DNA structures and minor oxidation of thiol linkages over extended storage periods. Critically, the sensor remained functional and within the calibrated linear range throughout the 21-day storage period, indicating that the tripodal thiol anchoring strategy provides sufficient long-term immobilization stability for practical use.

**Figure 8.**
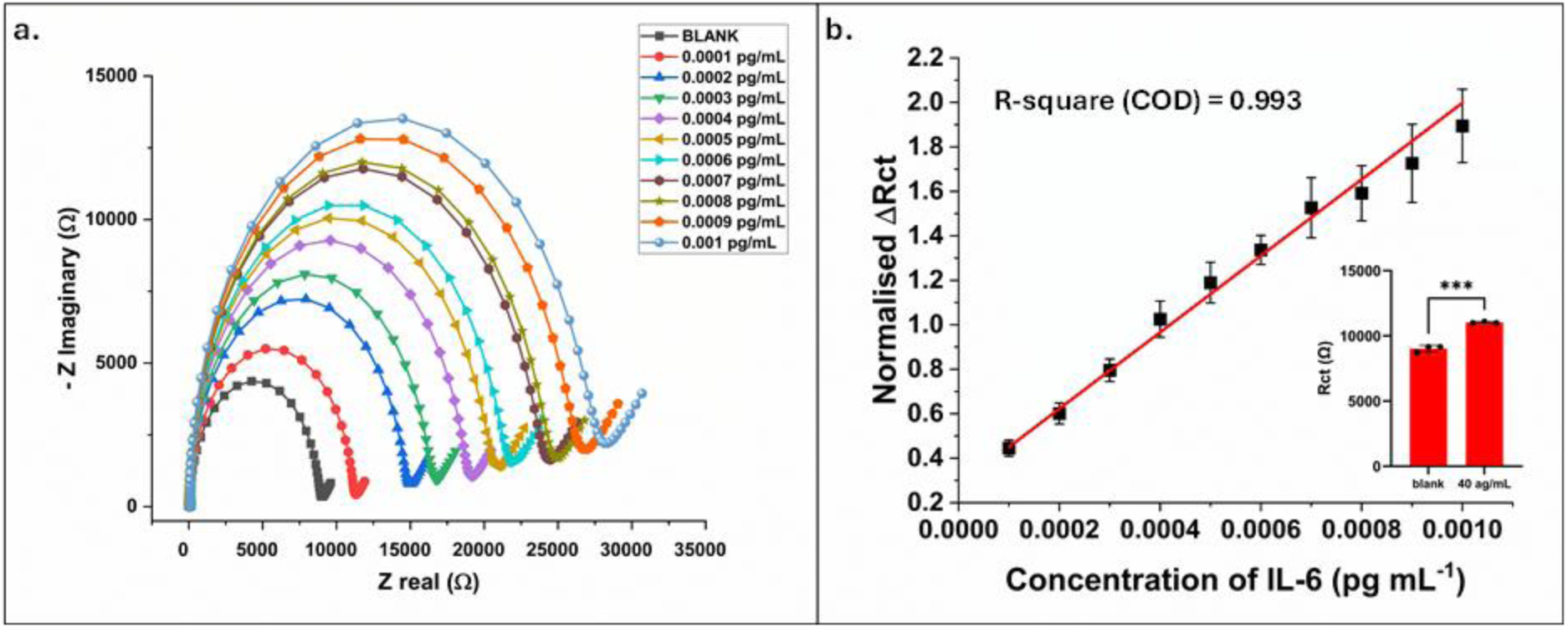
Analytical performance of the AP-S-TDN-Au-SPE impedimetric biosensor for IL-6 detection. (a) Representative Nyquist plots recorded after incubation with increasing concentrations of IL-6 (0.0001 to 0.001 pg mL⁻¹) in PBS (pH 7.4). Solid lines: fit to the Randles equivalent circuit. (b) Calibration curve plotting impedimetric response as a function of IL-6 concentration (0.0001 to 0.001 pg mL⁻¹). Linear fit: Normalised ΔRct = [1719.10592 ± 49.72008] (concentration of IL-6 in pg mL^-1^) + [0.27854 ± 0.01894], R² = [0.99335]. Linear range: 0.0001–0.001 pg mL⁻¹. LOD = 55 aM (3σ/slope criterion; σ = SD of the first lowest concentration, n = 3). Sensitivity = 1.55x 10^7^ Ω (pg mL⁻¹)⁻¹. All data shown as mean ± SD (n = 3 independent electrodes per concentration).

## 4. Conclusions

In this work, we have demonstrated a programmable impedimetric aptasensor platform built on tetrahedral DNA nanostructure (TDN)-engineered gold screen-printed electrodes (Au-SPEs) for the ultrasensitive and selective label-free detection of interleukin-6 (IL-6). The central contribution of this study is the systematic elucidation of how the number of thiolated base vertices — zero, one, two, or three — governs the interfacial charge-transfer resistance and, consequently, the analytical performance of the sensor. Across all configurations tested, a clear monotonic relationship was observed between multipodal anchoring geometry and baseline Rct, increasing from free oligonucleotide films (lowest Rct) through single- and double-vertex-thiolated TDNs to the fully tripodal AP-S-TDN architecture (highest Rct of 9010.97 ± 205.32 Ω). This hierarchy demonstrates unambiguously that thiolated vertex number is not merely a stability parameter but a primary determinant of interfacial film order, nanochannel geometry, and signal transduction efficiency — a design principle that, to our knowledge, has not been previously quantified in a systematic vertex-by-vertex study on disposable Au-SPEs. Mechanistic analysis through Randles equivalent-circuit modelling revealed that the exceptional sensing performance of the tripodal AP-S-TDN interface arises from the cooperative effects of three structural features: robust Au–S chemisorption at all three base vertices, which locks the tetrahedron upright and prevents tilt-induced conductance leakage; a rigid three-dimensional geometry that positions the anti-IL-6 aptamer at a precise apex projection into solution, maximising target accessibility; and a compact, well-ordered monolayer that confines the dominant electron-transfer pathways to narrow nanochannels between adjacent tetrahedra. Upon IL-6 binding, steric occlusion and additional charge at the apex aptamer efficiently block these channels, producing a disproportionately large concentration-dependent increase in Rct. This coupling between local molecular recognition events and global interfacial impedance changes underpins the steep calibration slope (1,719.1 ± 49.7 normalised ΔRct per pg mL⁻¹, R² = 0.993) and the exceptionally low limit of detection of 55 ag mL⁻¹ achieved by the final sensor. Under fully optimised conditions — pH 7.0, 0.05 µM TDN immobilisation concentration, and MCH passivation at 37 °C for 1.5 h — the AP-S-TDN-Au-SPE sensor exhibits a linear dynamic range of 0.0001 to 0.001 pg mL⁻¹ with a sensitivity of 1.55 × 10⁷ Ω (pg mL⁻¹)⁻¹. Selectivity evaluation against a panel of seven physiologically relevant interferents confirmed excellent specificity, with five of seven compounds — TNF-α, glucose, ascorbic acid, cysteine, and glycine — producing relative responses below 10% of the IL-6 signal, and no interferent exceeding 14.4%. The sensor demonstrated robust fabrication reproducibility with intra-electrode, inter-electrode, and batch-to-batch %RSD values of 2.8%, 4.37%, and 5.1% respectively, and retained 95% of its initial response after 21 days of refrigerated storage — confirming that the tripodal Au–S anchoring strategy provides sufficient long-term operational stability for practical deployment. Taken together, these findings establish tripodal thiolation of the TDN base as a critical and tunable design parameter for impedimetric aptasensors, providing a generalizable framework for engineering electrochemical interfaces with programmable charge-transfer characteristics. The use of disposable Au-SPEs as the substrate renders the platform compatible with portable potentiostat hardware, supporting future translation to point-of-care cytokine monitoring in sepsis management, oncology surveillance, and autoimmune disease stratification. Future work will focus on extending the calibration range to clinically prevalent IL-6 concentrations in the pg mL⁻¹ to ng mL⁻¹ regime, reducing the TDN immobilisation time for single-step point-of-care deployment, incorporating structural characterisation by TEM and AFM to directly visualise the interfacial TDN architecture, and expanding the platform to simultaneous multiplexed detection of cytokine panels relevant to systemic inflammatory syndromes.

## Notes

The authors declare that they have no known competing financial interests or personal relationships that could have appeared to influence the work reported in this paper.

## Acknowledgments

A.K.Y. acknowledges the financial support received from the Anusandhan National Research Foundation (ANRF) under the ANRF-National Postdoctoral Fellowship (PDF/2025/007589).

D.B. thanks SERB, GoI, for the Ramanujan Fellowship, DST-Nidhi Prayas for the start-up grant, and Gujcost-DST, GSBTM, BRNS-BARC, and HEFA-GoI, MoES, for STARS for research grants.

## CRediT authorship contribution statement

**Bhagyesh Parmar:** Methodology, Investigation, Formal Analysis, Data Curation, Writing – Original Draft. **Dhiraj Bhatia:** Funding acquisition, Project administration, Resources, Supervision, Validation, Writing – review & editing. All authors have read and approved the final version of the manuscript. **Amit K. Yadav:** Conceptualization, Methodology, Supervision, Writing – original draft, Project administration, Supervision, Validation, Visualization, Writing – Review and Editing.

## Data Availability Statement

No new data were generated or analyzed in this study.

## Declaration of Competing Interest

### Declaration of Generative AI and AI-assisted technologies in the writing process

During the preparation of this work, the author(s) used ChatGPT4 to improve readability and language. After using this tool/service, the author(s) reviewed and edited the content as needed and take(s) full responsibility for the content of the published article.

